# Superhydrophobic Rotation-Chip for Computer-Vision Identification of Drug-Resistant Bacteria

**DOI:** 10.1101/2023.01.06.523046

**Authors:** Jiacheng He, Ruonan Peng, Henry Yuqing, Rafi Karim, Juhong Chen, Guoyu Lu, Ke Du

**Affiliations:** Department of Mechanical Engineering, Rochester Institute of Technology, Rochester, NY, 14623 USA; Department of Chemical and Environmental Engineering, University of California, Riverside, 900 University Ave, Riverside, CA, 92521 USA; Department of Manufacturing and Mechanical Engineering Technology, Rochester Institute of Technology, Rochester, NY, 14623 USA; Department of Biomedical Engineering, Rochester Institute of Technology, Rochester, NY, 14623 USA; Department of Biological Systems Engineering, Virginia Tech, Blacksburg, VA, 24061 USA; School of Electrical and Computer Engineering, University of Georgia, Athens, Georgia 30602 USA

**Author notes:** Corresponding Author, Present Addresses 900 University Ave, Riverside, CA, 92521 USA.

**Keywords:** Microfluidics, superhydrophobicity, *Escherichia coli (E. coli)*, point-of-care (POC), beta-galactosidase, drug-resistance, computer vision

## Abstract

The transport, distribution, and mixing of microfluidics often require additional instruments, such as pumps and valves, which are not feasible when operated in point-of-care (POC) settings. Here, we present a simple microfluidic pathogen detection system known as Rotation-Chip that transfers the reagents between wells by manually rotating two concentric layers without using external instruments. The Rotation-Chip is fabricated by a simple computer numerical control (CNC) machining process and is capable of carrying out 60 multiplexed reactions with a simple 30-degree or 60-degree rotation. Leveraging superhydrophobic coating, a high fluid transport efficiency of 92.78% is achieved without observable leaking. Integrated with an intracellular fluorescent assay, an on-chip detection limit of 1.8×10^6^ CFU/mL is achieved for ampicillin-resistant *Escherichia coli (E. coli)*, which is similar to our off-chip results. We also develop a computer vision method to automatically distinguish positive and negative samples on the chip, showing 100% accuracy. Our Rotation-Chip is simple, low-cost, high-throughput, and can display test results with a single chip image, ideal for various multiplexing POC applications in resource-limited settings.

## INTRODUCTION

Food- and water-borne infections, including diarrhea, gastrointestinal ailments, and systemic illnesses, present a major health threat to individuals and healthcare facilities^1^. Many pathogens, such as *Escherichia coli* (*E. coli*), could lead to food- and water-borne illnesses and have been reported to cause considerable morbidity and mortality^2^. Early detection of bacteria not only increases the likelihood that patients receive appropriate medical treatment but also prevents the spread of these pathogens and reduces the risk of antimicrobial resistance^3,4^. The conventional method for bacteria detection is plate culturing^5^. Despite being relatively inexpensive and reliable, plate culturing is time-consuming, commonly 2 to 3 days^6^, and each positive culture is labor-intensive^7^. Moreover, injured or nonculturable microorganisms could result in insufficient detection capacity of plate culturing^8^. Polymerase chain reaction (PCR) is a widely used molecular-based method for bacterial detection^9^; however, PCR involves thermocyclers and sophisticated bulky equipment, which hardly make it readily conducive to Point-of-Care (POC) diagnostics^10^. Besides, it has limited multiplexing capability due to the design requirement for primer sets to be non-complementarity and to anneal at similar temperatures^11^. Typically, only three to four target sequences are detected simultaneously in a single reaction^12^. Therefore, multiplexed high-throughput POC detection, which could screen various analytes such as pathogens simultaneously, would be highly desirable because it allows for a rapid, affordable, and reliable qualitative readout.

Following the World Health Organization’s ASSURED Challenge (Affordable, Sensitive, Specific, User-friendly, Rapid and robust, Equipment-free, and Deliverable)^13^, recent research efforts have been focusing on culture-independent POC approaches. Economically, the market demands high throughput to maximize time and cost efficiency. Integrated biological systems like microfluidic chips can meet these criteria^6^. With a high degree of automation, many of these portable devices can analyze a large number of samples at once and output detection signals qualitatively^14^. Qualitative biosensors can reduce the cost of detection even more by not relying on sophisticated instruments such as a spectrometer^15^ or a smartphone-based microscope^14^. For example, three-dimensional microfluidic paper-based analytical devices (3D-μPADs) offer an affordable solution for analyzing multiplexed assays^16^. However, one major challenge with μPADs and all different kinds of microfluidics is distribution homogeneity, where channels must have the same hydrodynamic resistance for every branch in a certain region to prevent a bias in the final output. Additionally, the use of adhesive makes them irreversibly bound devices that cannot be reused^16^. For multiplexed detection on a microliter or smaller scale, microfluidic systems often require pumps and valves to transport the sample to many reaction wells for the reaction to occur^17,18^. The requirement for hydraulic control and gas-permeable materials can make on-chip reagent storage difficult and increase the cost and complexity of operation^19^.

A notable example of a qualitative microfluidic detection device is the SlipChip^19,20^. By using two pieces of etched soda-lime glass with chromium and photoresist coating, the SlipChip mechanically transports precise volumes of sample to mix with preloaded reagents without pumps or valves^19^. This device is designed to analyze very small volumes as it delivers samples in wells of only about 5 nL and a throughput of 48 individual reactions. However, evaporation prevention using a layer of fluorocarbon lubrication is necessary for the operation of the SlipChip, and reagent preloading has to be carefully carried out with volume-controlled droplets in specifically angled Teflon tubing. Mechanically, the device assembly lacks structural robustness since the two halves of the device are secured by nothing but two small binder clips. In addition, the SlipChip is fabricated via standard photolithography and glass wet etching, which is costly and requires a cleanroom setting^19^.

In this study, we present a microarray-based rotational microfluidic device designed for running high-throughput multiplexed pathogen detection assays. Known as the Rotation-Chip, this pocket-sized microfluidic system consists of two superhydrophobic coated polycarbonate layers that are fabricated by a computer numerical control (CNC) machining process. By simply manually rotating the disk, 60 individual samples are mixed with the reagents without causing inhomogeneity or leaking. To show an application of our chip, an intracellular fluorescent assay is used to detect ampicillin-resistant *E. coli*, demonstrating a detection limit similar to our off-chip results but does not require tedious manual pipetting. We also create a mean-shift algorithm to evaluate the fluorescence intensity of all the samples on the rotational chip, demonstrating 100% accuracy. The simple design of this chip solves the aforementioned problems of existing microfluidic devices by featuring a shearing-driven channel design for distribution homogeneity, a reversible and reusable assembly, superhydrophobic surface chemistry, and easy fabrication and operation processes, which is ideal for POC applications.

## MATERIALS AND METHODS

### Chip Fabrication

Measuring 80 mm in diameter, the top and bottom layers of the chip body were machined out of two pieces of 3-mm thick clear polycarbonate sheets by an automated CNC machine (ProtoTrak DPM5). On the one-step design variant, each well is 2.5 mm in diameter and 1 mm deep. The channels connecting the reagent array are of the same depth as the wells and 0.5 mm wide. Since a slight overfill is important for distribution homogeneity, these dimensions yield an effective fill volume of 5 μL per half well, 10 μL per assembled well, and 100 μL per assembled reagent array.

### Surface Treatment & Transport Efficiency

Surface treatments using two materials, Teflon (No. D14896684) and NeverWet^®^ (No. 274232), were conducted individually to evaluate and compare their hydrophobic performance. Before each surface treatment, the inner surface of each chip layer was cleaned in an ultrasonic bath of deionized (DI) water for 5 min and dried with an air gun. For surface treatment using Teflon, the Teflon solution was poured into a 100-mm Petri dish to form a pool about 3 mm deep. With its inner surface facing down, each layer was gently dipped into the Teflon solution until all features were covered. The layer was then lifted from the solution to let excess solution drip off while being slowly roasted by hand to achieve a reasonably even coating. After coating, both layers were placed, coated surface up, in their glass Petri dishes, and baked overnight at 80 □. The surface treatment using NeverWet^®^ was performed per the product manual. The “base coat” was sprayed onto the inner surface of each layer from approximately 20 cm away, then it was air-dried at room temperature for 30 min. The “top coat” was then applied in the same manner twice, 2 min apart. The coatings were allowed to cure at room temperature overnight before use.

### Assay and detection limit

Fluorescein di-ß-D-galactopyranoside (FDG) was purchased from Abcam. To prepare a 10 nM stock solution, 5 mg of FDG was dissolved in 761.6 μL of H_2_O: DMSO: EtOH (8:1:1) and stored at -20 □ in the dark. β-galactosidase (β-gal) powder (Sigma-Aldrich) with an activity of 1,000 U was dissolved in 1 mL of PBS. The solution was serially diluted by a factor of 10 into 0.001, 0.01, 0.1, and 1 U/mL for further use.

For the one-step assay, FDG was used to measure the intracellular activity of β-gal in LacZ strains, and ampicillin-resistant *E. coli* BL21 (DE3) (New England BioLabs, Inc.) with pUC19 plasmid (New England BioLabs, Inc.) was selected as the model stain. After the pUC19 vector was transformed into the *E. coli* strain by heat shock, LB-agar plates which contained 100 μg/mL ampicillin (Sigma-Aldrich) were used to grow *E. coli*. After the plates were incubated overnight at 37 □, a single colony of *E. coli* harboring pUC19 plasmid was then inoculated into LB broth media supplemented with 10 μg/mL ampicillin, followed by overnight incubation at 37 □ under 200 rpm. After that, 1 mL of the bacterial cells were pelleted in a centrifuge at 10,000 g for 5 min and resuspended in 1 mL of PBS. This step was then repeated twice. Next, the optical density (OD_600_) of the bacterial cells was measured to calculate the concentration, and serial dilution was performed in PBS for the harvested bacterial cells for further use.

Based on the volume of the one-step chip design, 5 μL of either β-gal or *E*.*coli* samples were added into 5 μL of FDG to perform off-chip experiments, and 5 μL of PBS was added instead as the negative group. The 10 μL mixture was vortexed and placed in a 37 □ water bath for 30 min under dark conditions to avoid the photobleaching of FDG. After the incubation, the mixture was excited by a blue light transilluminator (brand: SmartBlue™, Part number: NEB-E4100, excitation wavelength of 465 nm) for naked-eye observation. Finally, 60 μL of nuclease-free water (Thermofisher) was added to each tube, which was then characterized by a commercial spectrofluorometer (JASCO FP-8500, USA).

Prior to starting the on-chip experiment, the chip was sterilized with ultraviolet (UV) light for 30 min. Then, 100 μL of FDG was loaded into each reagent channel, and 5 μL of the sample was loaded into each sample well, along with 5 μL of PBS as a negative group. The top layer was slowly rotated 30° while holding the bottom layer steady; then the chip was placed into a Ziploc bag for waterproofing and incubated in a 37 □ water bath for 30 min in the dark. Finally, the rotated chip was placed onto the transilluminator to check the fluorescence.

### Image processing

Test sample circles are segmented by the mean-shift algorithm using intensity as the segmentation criteria^21^. Once the test sample circles are segmented, we apply Hough transform to detect each sample well and recognize the presence of the bacteria with various concentrations^22^.

## RESULTS AND DISCUSSION

**Figure 1a** shows the end of an *E. coli* detection assay, where qualitative detection results can be observed in a Rotation-Chip under the UV light of a transilluminator. **Figure 1b** shows the Rotation-Chip as an assembly of a pair of CNC machined polycarbonate disks with varying features. The top layer is designed to rotate clockwise relative to a static bottom layer about the central post. A guide pin and a guide rail system ensure the rotation is constrained to a fixed path with a physically defined initial position, final position, and rotational freedom of 30 degrees. An interference fit between the post and the central top layer hole ensures precise detuning and prevents the top layer from falling out of the assembly with ease. When assembled, the two layers of the one-step chip form 6 groups of radially symmetrical test groups, each capable of running 10 tests. Each test group contains a reagent array and a sample array positioned along the same rotational path. The reagent array contains 10 reagent wells of 10 μL whose bottom halves are connected by a continuous channel on the bottom layer. The channel inlet to the left of the channel is designed to fit most micropipette tip sizes during loading, and the channel outlet is through a small hole in the top layer for pressure equalization. Together, these features enable the user to fill all regent wells at once regardless of the number of wells in the array, avoiding repetitive pipetting and saving operation time. On the other hand, the sample array contains wells of the same number and size as the reagent array but is kept separate to avoid contamination across different tests. To make sure the assembly process takes place before the loading process, the top halves of the sample wells are through holes so the user can individually load samples post-assembly. **Figure 1c** and its accessory cross-sectional views show the operation mechanics of running a test group in an assay, where the blue and the red fluids represent the reagent and sample, respectively. To start the fluid transportation process, the user fills both layers of the reagent array with the reagent before rotating the top layer while holding the bottom layer still.

**Figure 1.**
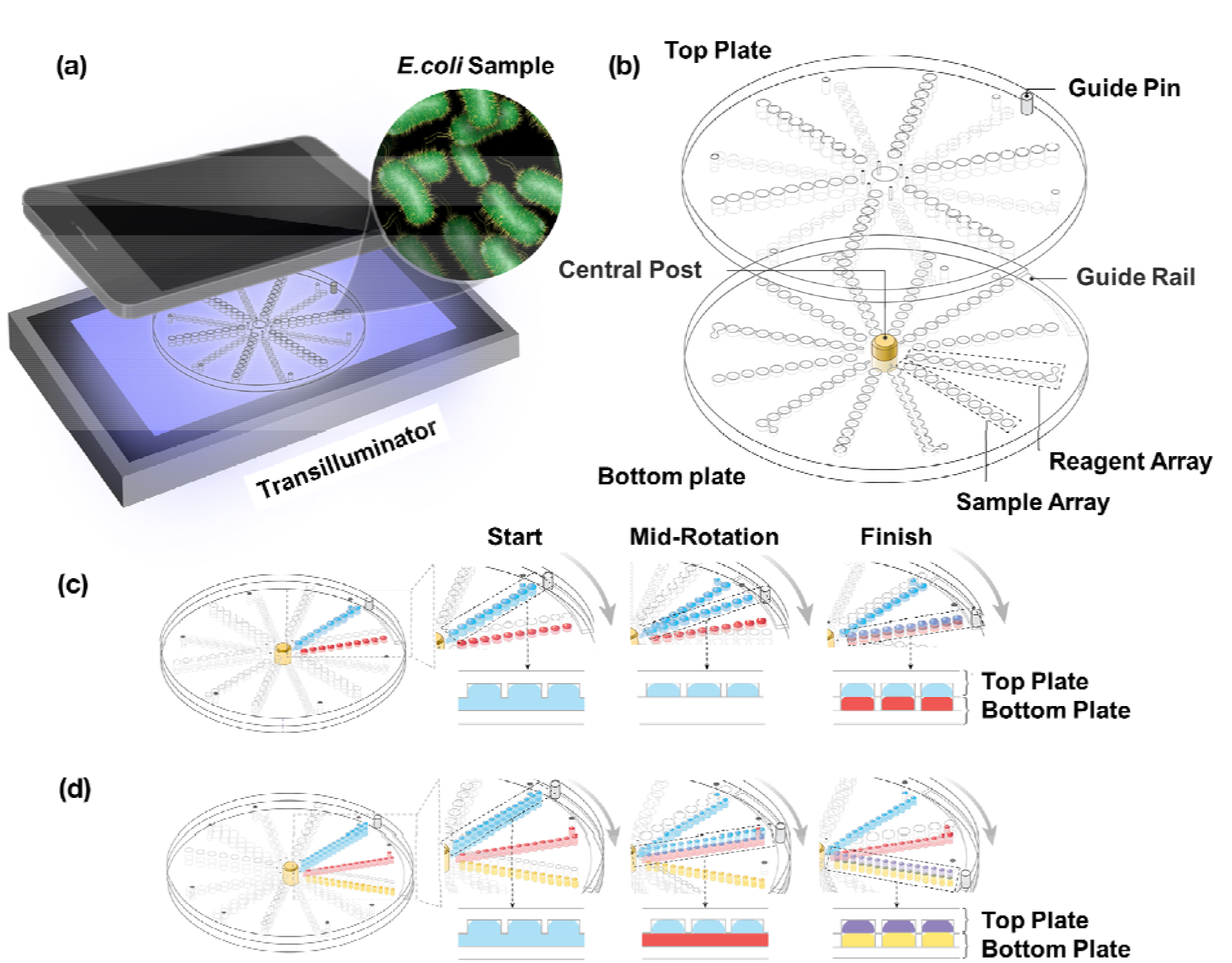
Rotation-Chip design and operation. (a) Fluorescence is excited by a transilluminator and the results are read by either naked dye or a smartphone. (b) Exploded view showing the mechanical components of the one-step chip design. Operational schematic of the fluid transportation of the (c) one-step chip design and (d) two-step chip design.

During the rotation, this shearing motion carries discrete volumes of the reagent fluid and carries them along the rotational path. The rotation operation is complete when the guide pin touches the end of the guide rail as the reagent fluid makes contact with the sample in each sample well. In **Figure 1d**, the two-step chip executes the same fluid dynamics, but the addition of the secondary reagent produces a mixture of the two reagents that mixes the sample at the end of a 60-degree rotation, shown in red, purple, and yellow, respectively.

**Figure 2a** shows the contact angles of Rotation-Chip with no coating, Teflon coating, and NeverWet^®^ coating, respectively (from left to right). Among them, NeverWet^®^ coating processes the most hydrophobic property, of which the static contact angle is 156.8 °. The SEM images of the Rotation-Chip before and after the NeverWet^®^ coating are shown in **Figure 2b**. The NeverWet^®^ coating consists of fluorocarbon-coated silica particles to produce surface roughness and native superhydrophobicity, as shown in the zoom-in view, while the uncoated chip shows a clean surface. To measure the number of wells leaked during the rotation process and to determine the coating on which side of the chip would provide the best transport efficiency, food dye was used as a model for reagents and samples. After the food dye was loaded on the Rotation-Chip, the chip was rotated 30° clockwise and the number of leaked wells was counted by the naked eye. The experiment for each coating situation was repeated 5 times to minimize the error. The leak count for the NeverWet^®^ coated Rotation-Chip under different conditions is shown in **Figure 2c**. No coating was represented by ‘-’ while NeverWet^®^ coating was represented by ‘+’. The first symbol represented the coating situation for the top layer, and the second symbol represented the coating situation for the bottom layer. When both layers were coated with NeverWet^®^, the leakage of Rotation-Chip was significantly minimized. The transport efficiency of the Rotation-Chip was tested with a dedicated chip for testing, where there was only one well on the top layer of the Rotation-Chip. The weight of 10 µL of the food dye was measured with the high precision laboratory balance (Intelligent PM-300), then 10 µL of the food dye was added into the well and rotated for 5 circles. After that, the remaining food dye in the well was removed by a pipette, which was then weighed with the same balance to calculate the transport efficiency of the Rotation-Chip. The experiment for each coating situation was repeated 5 times to minimize the error. **Figure 2d** shows the transport efficiency results for uncoated, Teflon-coated, and NeverWet^®^ coated Rotation-Chip. The NeverWet^®^ coated chip transports 92.78% of the reagent while that of Teflon coating is only 86.02%. For the chip without any coating, all the food dye leaked out of the wells during the rotation, resulting in a transport efficiency of 0. **Figure 2e** is the food dye demonstration on the two-step chip before and after the NeverWet^®^ coating. Before the NeverWet^®^ coating (top row), there was liquid leakage during both reagent loading and chip rotation. After the NeverWet^®^ coating (bottom row), the liquid leakage was significantly minimized, and this avoided sample contamination and improved the reagent utilization, which provided a more accurate results readout. Also, the variety of well sizes demonstrates the morphological multiplexing ability of a two-step chip. With appropriate well dimension adjustment, the Rotation-Chip may be easily compatible with some two-step assays that require multiple mixing steps.

**Figure 2.**
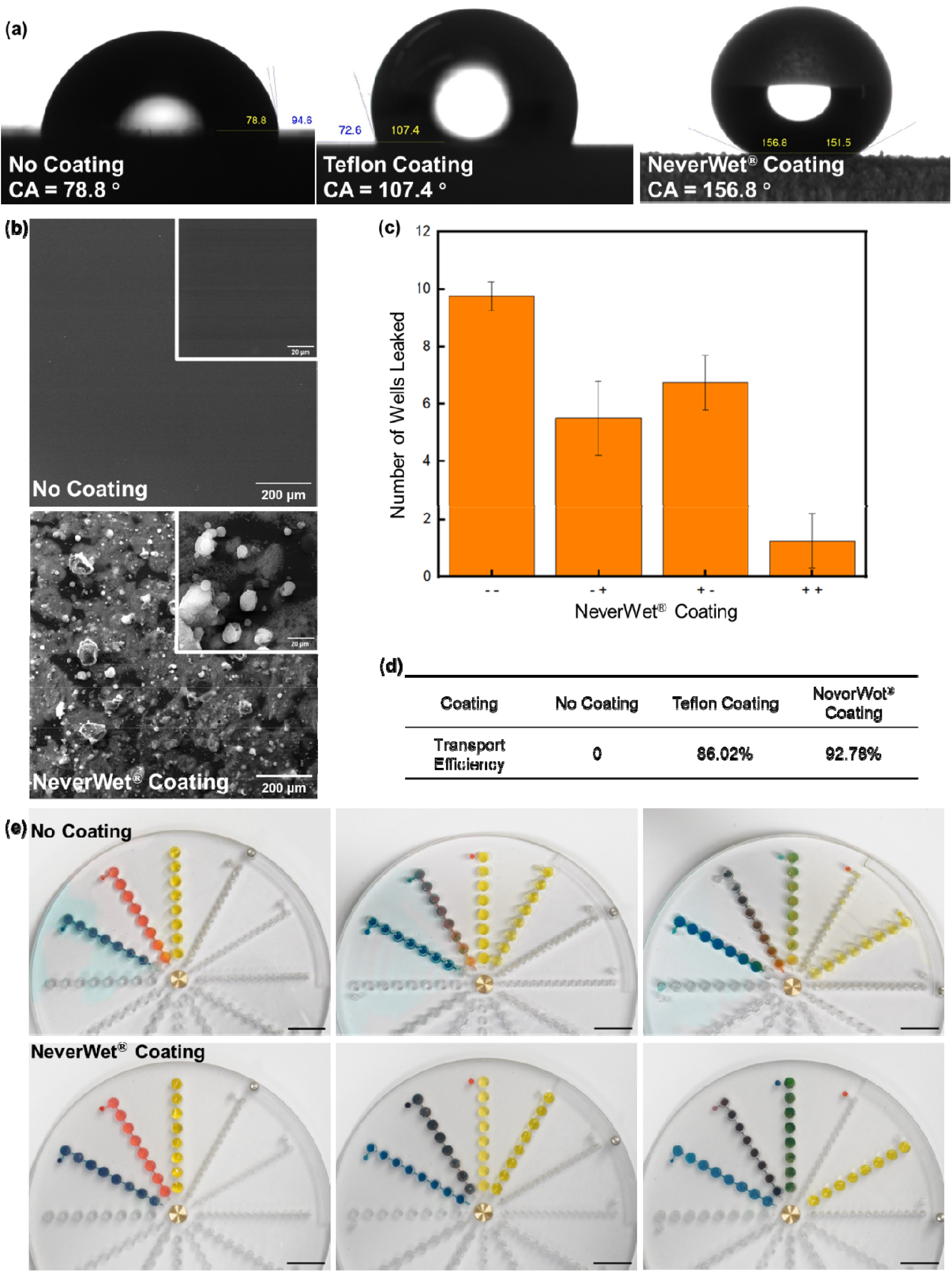
Surface treatment and transport efficiency of Rotation-Chip. (a) The static contact angles of Rotation-Chip with no coating, Teflon coating, and NeverWet^®^ coating (from left to right). (b) SEM images of Rotation-Chip before and after the NeverWet^®^ coating. (c) Rotation-Chip leak count for NeverWet^®^ coatings under different conditions. No coating is represented by ‘-’ while NeverWet^®^ coating is represented by ‘+’. The first symbol represents the coating situation for the top layer, and the second symbol represents the coating situation for the bottom layer. (d) Transport efficiency of the Rotation-Chip with Teflon and NeverWet^®^ coating. (e) The food dye demonstration on the two-step chip before and after the NeverWet^®^ coating (scale bar: 10 mm).

*Escherichia coli* (*E. coli*) can produce the enzyme β-galactosidase (β-gal) to break lactose into galactose and glucose^23^. Fluorescein di-ß-D-galactopyranoside (FDG) is one of the most sensitive and versatile fluorogenic substrates for the β-gal^24^. **Figure 3a** shows the two-step sequential hydrolysis of non-fluorescent FDG by β-gal, first converting it to mono-beta-D-galactosidase (FMG) and then to highly fluorescent fluorescein (Excitation/Emission, 488/512), which can be detected by an LED transilluminator^25^. The fluorescence intensity (512 nm) of off-chip results of the assay in response to purified β-gal with different activities (0 - 1 U) is shown in **Figure 3b**. After a 30 min reaction, the fluorescence intensity was evaluated by a commercial spectrofluorometer (JASCO FP-8500, USA). As shown in **Figure 3c**, more fluorescein is produced by FDG hydrolysis with the increasing β-gal concentration, and the fluorescence intensity is increased. For the on-chip experiment, 100 µL of FDG was added to the reagent array, and 5 µL of β-gal was added sequentially in each well. After rotation and incubation, the chip was excited from the bottom with the LED illuminator, and the results are shown in **Figure 3d**. When the β-gal activity is greater than 0.01 U, the fluorescence differences between positive and negative groups can be observed on the transilluminator by the naked eye. This detection sensitivity is comparable to the off-chip results.

**Figure 3.**
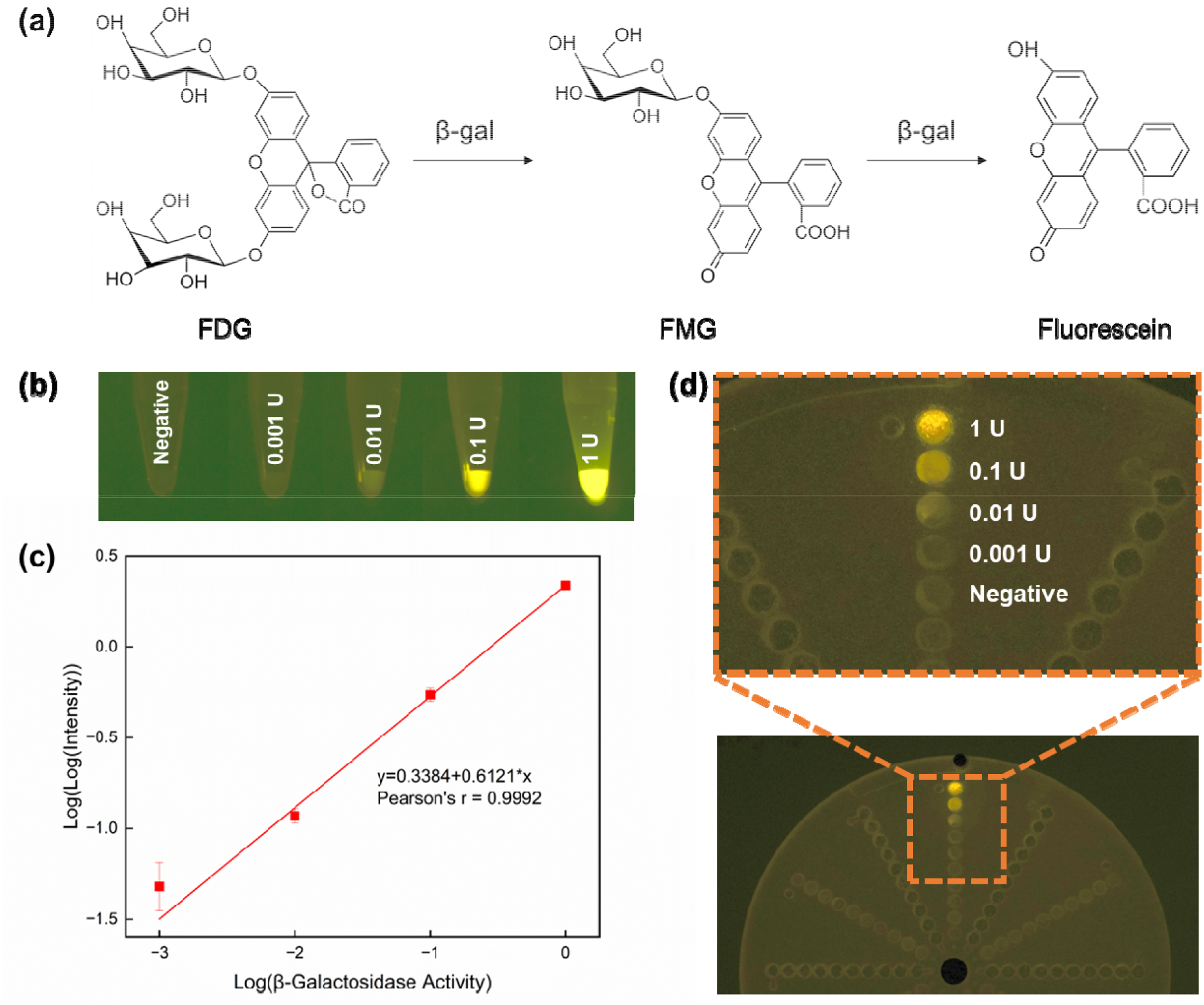
(a) Schematic of β-gal sequential hydrolyzation of non-fluorescent FDG. (b) Fluorescence intensity (512 nm) of off-chip results in response to β-gal with different activities (from 0 - 1 U). (c) Off-chip results of fluorescence intensity (512 nm) correspond to various concentrations of the β-gal. The linear region of the Log versus (Log (fluorescence intensity at 512 nm)) plotted as a function of the logarithmic concentration of β-gal (from 0.001 - 1 U) is highlighted (Pearson’s r = 0.9992). (d) On-chip detection results of β-gal (from 0 - 1 U) on One-Step Chip coated with NeverWet^®^.

We then optimized the *E. coli* detection by varying the FDG concentration ranging from 0.01 to 0.25 nM. As shown in **Figure 4a**, when *E*.*coli* concentration is higher than 1.8 × 10^6^ CFU/mL, the fluorescence differences between positive and negative groups can be easily observed by the naked eye. The fluorescence intensity of each group was then quantified by the Spectrofluorometer, and the results at an FDG concentration of 0.25 nM are shown in **Figure 4b**. When the FDG concentration is 0.25 nM, the integrated fluorescence signal is highest with the best log-log linear fitting (Pearson’s r = 0.9734) among the four groups, as shown in **Figure 4c**. Thus, we selected 0.25 nM of the FDG for on-chip experiments. The FDG and *E*.*coli* loaded chip before incubation is shown in **Figure 4d** (top), with *E*.*coli* concentration ranging from 1.8 × 10^6^ to 9 × 10^7^ CFU/mL. When the *E*.*coli* concentration is higher than 1.8 × 10^6^ CFU/mL, the fluorescence signal is easily observable, as shown in **Figure 4d** (bottom), which is comparable to the off-chip results. In addition, three independent sets of the on-chip experiments (a total of 12 samples) were performed simultaneously by a single rotation, and the fluorescence readings are consistent among the three sets.

**Figure 4.**
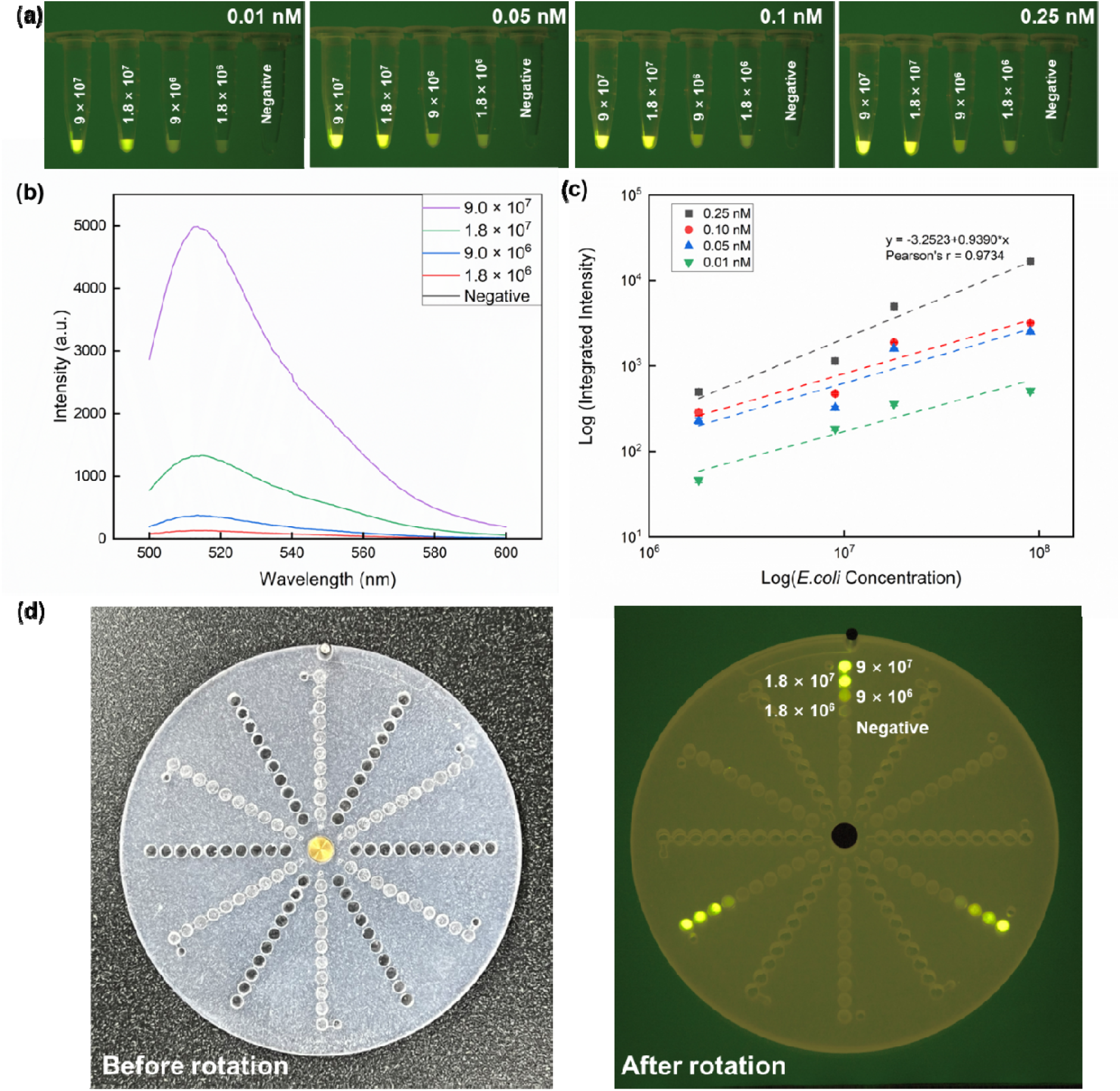
(a) Off-chip detection of *E. coli* (1.8 × 10^6^ - 9 × 10^7^ CFU/mL) with FDG of 0.01, 0.05, 0.1, and 0.25 nM (from top to bottom). (b) Off-chip results of fluorescence intensity corresponding to various concentrations of the *E. coli* (1.8 × 10^6^ - 9 × 10^7^ CFU/mL) at an FDG concentration of 0.25 nM. (c) Off-chip results of integrated fluorescence intensity (560 - 580 nm) corresponding to various concentrations of the *E. coli* (1.8 × 10^6^ - 9 × 10^7^ CFU/mL) on a logarithmic scale (Pearson’s r = 0.9734). (d) Detection of *E. coli* at different concentrations (1.8 × 10^6^ - 9 × 10^7^ CFU/mL) with 0.25 nM FDG on One-Step Chip coated with NeverWet^®^.

On-chip automatic recognition can be considered as a binary classification problem^26^, either positive or negative. As fluorescence intensity presents the major difference between positive and negative samples, we extract the color intensity histogram feature to represent each segmented sample and classify the samples based on each sample’s color intensity histogram feature^27^. To remove the background influence, 0 values in the histogram are removed, as 0 represents the black color, which can increase the difference between positive and negative color intensity histogram features. Due to the limited number of training samples and the nature of the binary classification tasks, we apply a decision tree to classify the testing^28^. The entire pipeline for processing images and classifying the testing samples is shown in **Figure 5**. Eventually, we achieved 100% classification accuracy to separate the detected positive samples from the negative samples.

**Figure 5.**
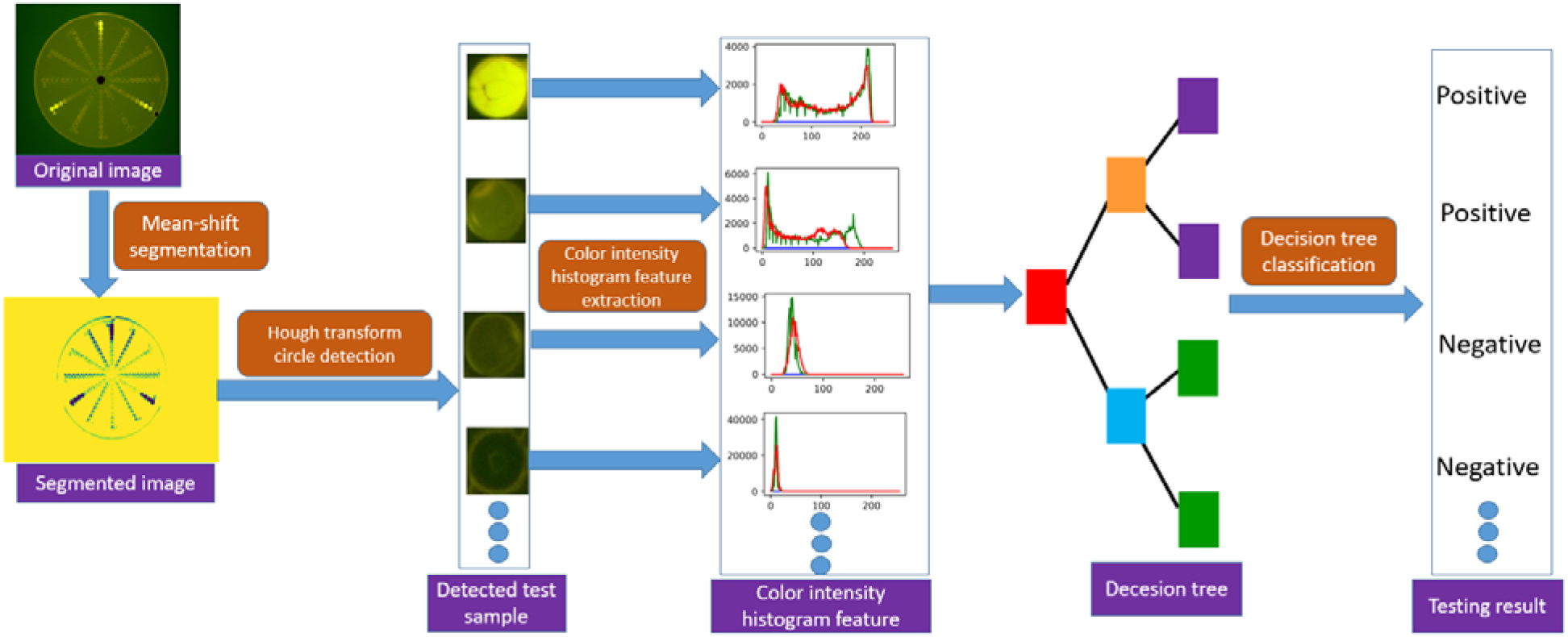
The image processing and testing result classification pipeline of computer vision enabled *E. coli* sensing on the superhydrophobic Rotation-Chip.

Due to its design, operation, and manufacturability, the Rotation-Chip has many advantages compared to other POC systems for pathogen detection. Firstly, the design of the Rotation-Chip results in good portability, throughput, and reliability. The motion in which a top plate slides across a bottom plate has been the traditional method to mechanically manipulate fluids through wells and channels. However, this linear motion disrupts the overall shape of the device and limits throughput per unit area by extending the top plate outside the boundary of the preoperative chip, exposing inner features or treated surfaces like in the case of the SlipChip^19,20^. By adopting a rotational design, the Rotation-Chip can deliver the same fluidic manipulation while keeping all sensitive features in a covered state and staying mechanically intact. Such a design lowers the chip’s spatial footprint and increases its maximum feature density for more portability and throughput. To ensure a reasonably easy rotation effort, the top and bottom plates of the Rotation-Chip are intentionally unclamped, with the assembly held together by only gravity and the friction from the mating between the central post and the central top plate hole. This design makes leakage prevention a crucial challenge to overcome for the fluid transport process to work with meaningful transport efficiency. Specifically, we found that surface treatment for hydrophobicity to be an effective leakage prevention solution without compromising mechanical integrity or needing dedicated sealing. Notably, we can achieve a superhydrophobic surface by using a commercially available spray coating, which is inexpensive and easy to apply. The low surface energy from the superhydrophobicity contributes positively to the Rotation-Chip’s ability to transport discrete volumes of fluid efficiently by keeping the intra-droplet surface tension high enough to withstand the shear stress from the rotation and minimize inter-plate leaking.

Secondly, the simplistic design of the Rotation-Chip results in an easy manual operation of a simple four-step process: load, rotate, incubate, and read. The user can follow the protocol described above and obtain a qualitative result without any expertise in using complicated lab equipment or interpreting fluorescence curves. In addition, all 120 wells can be screened at the same time with our computer vision method, making our Rotation-Chip a powerful POC system. Working together with a simple two-part assembly, the superhydrophobic surfaces of the Rotation-Chip allow for easy cleaning and reusability of at least five times as of this study. Another way to reduce operational complexity and cost is through system integration. Thanks to its constant size, the bottom of the Rotation-Chip can be equipped with a custom micro hot plate that can be configured to the temperature requirement of the assay, inexpensive and plug-and-play^29^. This reduces process complexity and further decreases cost by removing the need to use a water bath and its accessory supply, such as a sealable bag, to keep water from entering the chip during incubation.

Lastly, several popular methods of microfluidic chip fabrication are available to produce the Rotation-Chip, including polydimethylsiloxane (PDMS) molding^30^, glass etching^31^, CNC machining^32^, and injection molding^33^. Though conventional and inexpensive, PDMS molding has properties that were unsuitable for the shear strength and overall rigidity requirements of our application. For the prototypes shown in this study, CNC machining was chosen to be our method of fabrication. Compared to lithographic methods such as wet or dry etching, which are done on brittle material and are time-consuming^33^, CNC machining has a much faster lead time and facilitates the dimensional customizability of the design by allowing for the agile production of several design variants. A downside of CNC machining is the surface finish. Since the machined surface is always the inner functional surface in subtractive manufacturing, micro-marring caused by the CNC tool was unavoidable. This poor surface finish increased the surface shear stress, which contributed negatively to the transport efficiency of the chip, as indicated by a decent but improvable experimental value of 92.78%. Another drawback of CNC machining lies in how small the chip features could be made, an uncommon constraint in microfluidic device manufacturing. For future applications where better surface finish and more unit-area throughput (maximum number of wells in a certain chip area reasonably visible to the naked eye) are needed, high-resolution 3D printing^34^ and injection molding^33^ can be good alternatives to the aforementioned manufacturing methods for future production of the Rotation-Chip. For example, injection molding could be an efficient and cost-effective alternative for large-scale fabrication. Using the same clear polycarbonate material or cheaper substitutes like acrylic, an injection molded chip could accommodate smaller microfluidic features to even the nanoliter scale and provide a smoother surface finish^19^ for better unit-area throughput and transport efficiency.

While the commercial FDG-β-gal assay applied to Rotation-Chip is lysis-free and sensitive, various other qualitative intracellular assays with even higher sensitivity comparable to PCR can be incorporated into the Rotation-Chip^35^. For example, by exploiting the redox pathway of *E. coli*, p-benzoquinone was used to mediate a sensitive and economical bioassay to detect *E. coli* concentrations as low as 1.0 × 10^4^ CFU/mL by providing a colorimetric readout in 1 hour^36^. Su et al. also developed a sensitive colorimetric method for *E. coli* detection that can be applied to the two-step chip with appropriate dimension customization. Mercaptoethylamine (MEA) could bind to *E. coli* through electrostatic adhesion, and it is also easily conjugated to gold nanoparticles (AuNPs) via the -SH group. When presented with *E. coli* under a certain pH condition, the MEA-AuNPs complex leads to a color change from blue to red in less than 5 min, which is observed by the naked eye^37^. Moreover, the chip can be employed for antimicrobial susceptibility testing assays^38^. These assays were developed to rapidly detect drug resistance caused by β-lactamase to aid in the treatment of urinary tract infections, which provides a sensitivity of 90.9% and specificity of 97.6% within a timeframe of 30 min. The caged enzyme amplifier (papain) is combined with a β-lactamase-targeting probe. When exposed to β-lactamase, the papain is activated, and this could further produce a colorimetric signal output via the hydrolysis of Nα-Benzoyl-L-arginine 4-nitroanilide hydrochloride (BAPA)^39^.

Besides healthcare, the Rotation-Chip has applications in a variety of other important areas, including food quality and environmental safety^40^. Due to its ability to deliver portable, equipment-free, affordable, user-friendly, rapid, and robust detection, the Rotation-Chip can be especially valuable in resource-poor environments, emergency situations, or at-home healthcare settings^41^. It can be coupled with isothermal or thermocycling amplification of nucleic acids^42^, stochastic confinement^43^, and reagent storage^19^. Due to its portability, it also makes a good candidate for testing environmental pollution from ambient samples^44^. In the future, a smartphone app can be developed by using our robust computer vision method and directly working with the Rotation-Chip for accurate, automatic, and high-throughput detection applications^45^.

## GRAPHICAL ABSTRACT

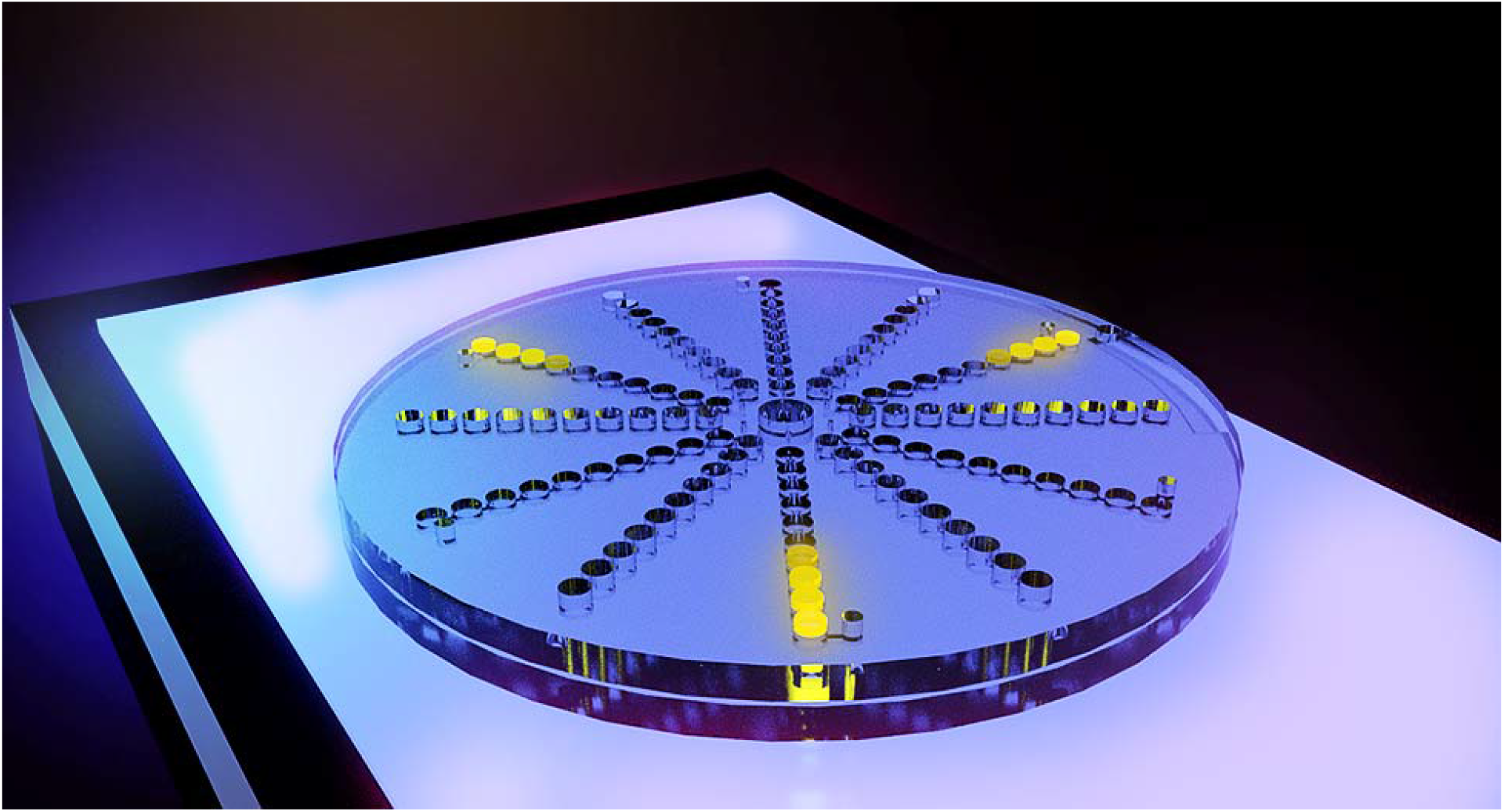

## Author Contributions

Jiacheng He, Juhong Chen, and Ke Du designed the experiments. Jiacheng He, Ruonan Peng, Henry Yuqing, and Rafi Karim conducted the experiments. Jiacheng He, Ruonan Peng, Guoyu Lu, and Ke Du wrote the manuscript. All the authors commented on the manuscript.

## ACKNOWLEDGMENT

The authors would like to thank Mary Nguyen for the schematic design.

## ABBREVIATIONS

FDG: Fluorescein-di-beta-D-galactopyranoside, β gal - β-Galactosidase

